# The representation of colored objects in macaque color patches

**DOI:** 10.1101/205104

**Authors:** Le Chang, Pinglei Bao, Doris Y. Tsao

## Abstract

An important question about color vision is: how does the brain represent the color of an object? The recent discovery of “color patches” in macaque inferotemporal (IT) cortex, the part of brain responsible for object recognition, makes this problem experimentally tractable. Here we record neurons in three color patches, middle color patch CLC (central lateral color patch), and two anterior color patches ALC (anterior lateral color patch) and AMC (anterior medial color patch), while presenting images of objects systematically varied in hue. We found that all three patches contain high concentrations of hue-selective cells, and the three patches use distinct computational strategies to represent colored objects: while all three patches multiplex hue and shape information, shape-invariant hue information is much stronger in anterior color patches ALC/AMC than CLC; furthermore, hue and object shape specifically for primate faces/bodies are over-represented in AMC but not in the other two patches.

## Introduction

We see the world in color because different objects are composed of materials with different reflectance spectra. The perception of color involves processing at multiple stages of the visual system. Recordings in early parts of the visual system reveal double-opponent cells in area V1 ^1,2^ and hue-selective cells in areas V2 and V4 ^3,4^; these cells allow the brain to compute the local hue at each location across a surface. But color processing does not end with extraction of local hue: the brain needs to integrate information about hue distributions across space with information about large-scale object shapes, to enable an organism to recognize and respond appropriately to colored objects. For example, in Fig. 1a, we readily perceive a red apple, which requires (1) correctly discriminating the object from the background, and (2) extracting the dominant hue within the object (Fig. 1a).

**Figure 1.**
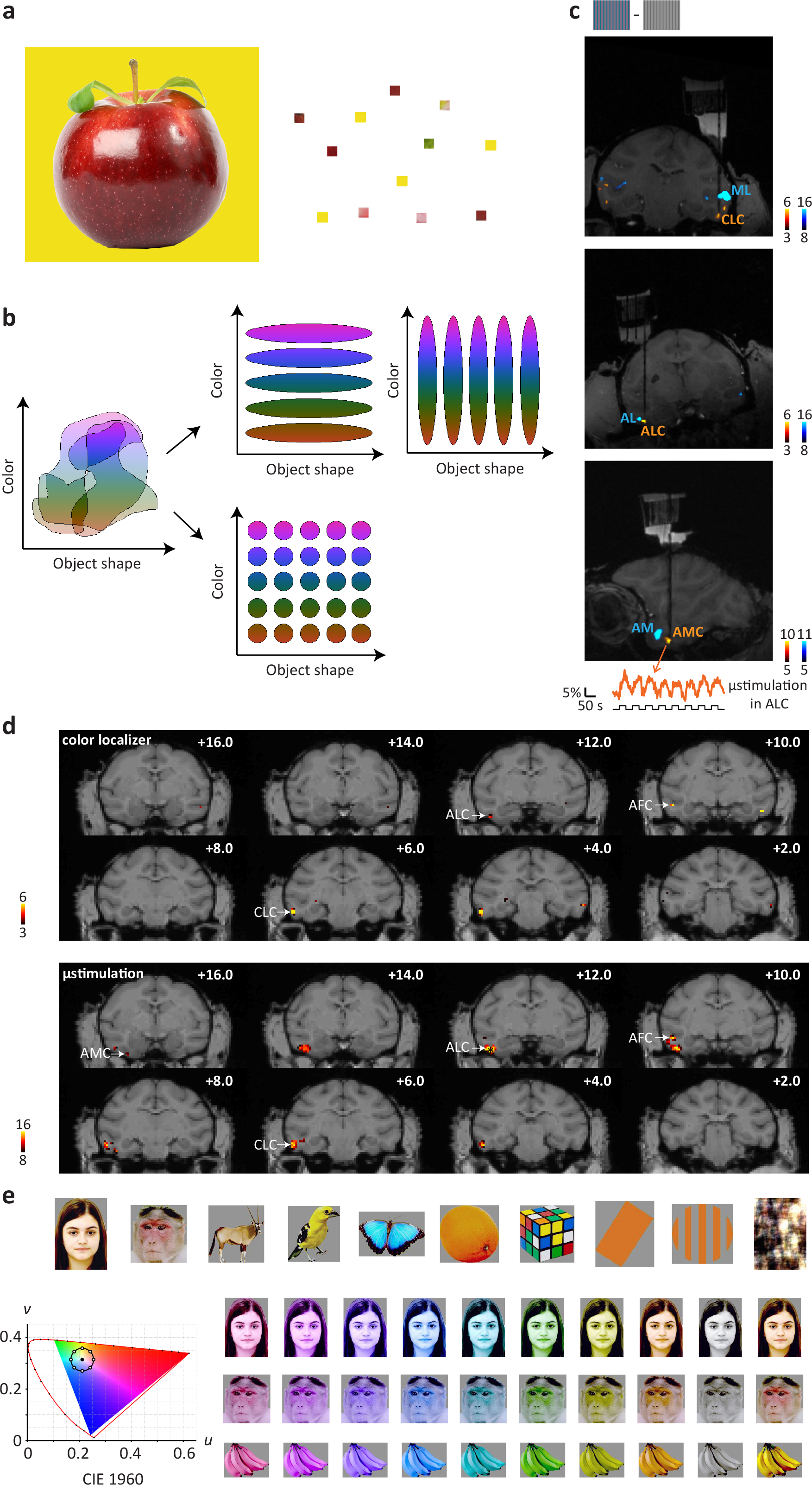
Recording sites, connectivity, and color stimuli. **a,** To identify the correct color of an object, e.g. an apple (*left*), local hue information (*right*) needs to be integrated with global shape information. **b,** Schemes for co-representing color and object shape information in visual system. Initially, before the visual system has explicitly segmented objects, color information and object shape information are largely entangled, with individual cells participating in the coding of multiple hues and object shapes. Two main strategies could be used to represent colored-objects in an organized way: 1) segregation of color and object information into parallel channels, resulting in object shape-invariant color-selective units and color-invariant shape selective units (top); 2) formation of units sharply tuned to both color and object shape (bottom). **c,** Coronal and Sagittal slices showing location of fMRI-identified face (blue) and color patches (yellow) in one monkey (M2) targeted for recording; dark black line indicates electrode. The most anterior color patch was not observed with fMRI using the color localizer in this animal, and was located by electrical microstimulation in ALC (bottom panel, changes in BOLD signal of the identified voxels during microstimulation shown below). Source: Tsao lab. **d,** Comparison between color patches identified by color localizer (top) and by microstimulation (bottom). The contrasts are overlaid on high-resolution coronal slices. Asterisk (*) indicates the stimulation site (ALC). The anterior-posterior position of each slice in mm relative to the interaural line is given in the top right corner. **e,** 82 images of 10 categories were used (see Supplementary Fig. 2a for all the stimuli). Each image underwent a series of transformation in hues. For each pixel of the image, luminance was kept constant, while chromatic coordinates (CIE 1960) fell on a circle with the same distance to “white” (filled circle) as the original pixel. Eight hues with different angles were used (open circles, starting from 0°, going clockwise at 45° step). A grayscale image with the same luminance and the natural color image were also presented.

In theory, there are two possible mechanisms by which the brain could effectively organize information about the color and shape of colored objects (we use the term “shape” to refer to all aspects of an object’s identity independent of hue, not simply overall shape). Color and shape information could be segregated into parallel channels, resulting in shape-invariant color-selective units and color-invariant shape-selective units (Fig. 1b, top). Alternatively, units could be sharply tuned to both color and shape (Fig. 1b, bottom). Our ability to name colors independent of shape (e.g., a “red” traffic light or a “red” apple) supports the first scheme, while our ability to respond to specific color-shape combinations (e.g., stop at a red traffic light, eat a red apple) supports the second scheme (though it’s also possible such semantic representations are not directly related to visual representations). In addition to asking how single cells encode shape and color at different stages of visual processing, we can also ask how the representation of colored objects at different stages of visual processing is transformed at the population level: are shape and color co-represented within intermingled cell populations along the entire visual pathway, or do they become segregated into separate populations?

To clarify how the representation of object color is transformed in the visual system following the extraction of local hue information at both the single cell and population level, we targeted fMRI-identified “color patches” in inferotemporal (IT) cortex ^5^ for electrophysiological recordings in three macaque monkeys. IT cortex has long been believed to be responsible for the representation of object shape ^6,7^, but a recent fMRI study revealed a set of regions in macaque IT selective for colored compared to grayscale gratings ^5^. Interestingly, color patches were yoked in position to face patches ^5^, regions in IT cortex selective for faces ^8^. The localization of color patches within IT cortex implicates their role in representing color within the context of object recognition, while the stereotyped relative localization of color patches and face patches raises the possibility that a hierarchical functional organization for processing colored objects exists, mirroring that for processing facial identity in the face patches ^9^. Previous studies of color processing in monkeys have mostly used artificial stimuli, such as white noise, sinusoidal gratings or simple geometric shapes ^5,10–12,^ while previous studies of object representation in IT have mostly used grayscale images, ignoring color variations ^13,14^. This is unsatisfying, given that *colored objects* are the only natural visual inputs. Here, we explored the co-representation of color and object identity by presenting images of objects systematically varied in color, while recording from color patches in IT.

## Results

### Recording sites, connectivity and stimulus generation

We first localized color patches in three monkeys with fMRI using colored versus black-white gratings ^5^ (Fig. 1c). This revealed the central lateral, anterior lateral, and anterior medial color patches (CLC, ALC, and AMC) in monkey M1, CLC, ALC, and the anterior fundus color patch (AFC) in monkey M2 (we could not find AMC in this animal), and CLC, ALC, and AFC in monkey M3 (we could not find AMC in this animal). Next, we electrically microstimulated color patches while performing simultaneous fMRI (see Methods), to reveal the anatomical connectivity of color patches, and to potentially identify AMC in monkeys M2 and M3. This technique has previously been used to study the connectivity of face patches ^15^, and to reveal a place-selective region downstream of another place-selective region that had been identified using fMRI ^16^. Stimulating ALC in monkey M2’s left hemisphere activated three additional patches. Two of these patches overlapped with AFC and CLC identified by the color localizer (Fig. 1d), while one of these patches was located anterior to the stimulation site, on the ventral surface of the inferotemporal gyrus medial to the anterior middle temporal sulcus. Based on its location and connectivity to ALC, we designated this patch AMC (Fig. 1c, d). Possibly this patch was actually missing and we found another one, however, importantly, physiology was consistent between monkeys for AMC identified in the two ways. Furthermore, stimulating CLC in the right hemisphere of monkey M2 activated a patch overlapping with ALC as identified by the color localizer (Supplementary Fig. 1). Overall, these results suggest that color patches in IT cortex form a strongly and specifically interconnected network, similar to face patches ^15^, and allowed us to identify a color patch anterior to ALC in an animal in which it was missing based on the color localizer experiments.

We next targeted middle color patch CLC, and anterior color patches ALC and AMC for electrophysiological recordings. For comparison, we also targeted several sites in IT outside the color patches, including face patch AM. To study the co-representation of color and object shape, we generated a stimulus set of 82 images from 10 different categories (Supplementary Fig. 2a), each rendered in 8 different hues (Fig. 1e; see Methods). In this way, we varied hue information and object shape information independently and simultaneously. Grayscale images and the original color images were also included in the stimulus set.

### Representation of hue by neurons in color patches

Color stimuli were presented for 200 ms (ON period) interleaved by a gray screen for 200 ms (OFF period) during recording. The full stimulus set was presented 7-10 times for each cell recorded. Responses of all cells in each patch as well as cells outside the color patches to grayscale images and a subset of color images are shown in Fig. 2. ANOVA analysis was performed to test the significance of color tuning (see Methods). 74.7% (65/87) of cells recorded in CLC, 84.6% (55/65) of cells in ALC and 88.9% (80/90) of cells in AMC were significantly color tuned (ANOVA, p<0.001); outside the color patches, only 27.3% (9/33) of cells were significantly color tuned. Within color patches, only significantly tuned cells were used for further analysis.

**Figure 2.**
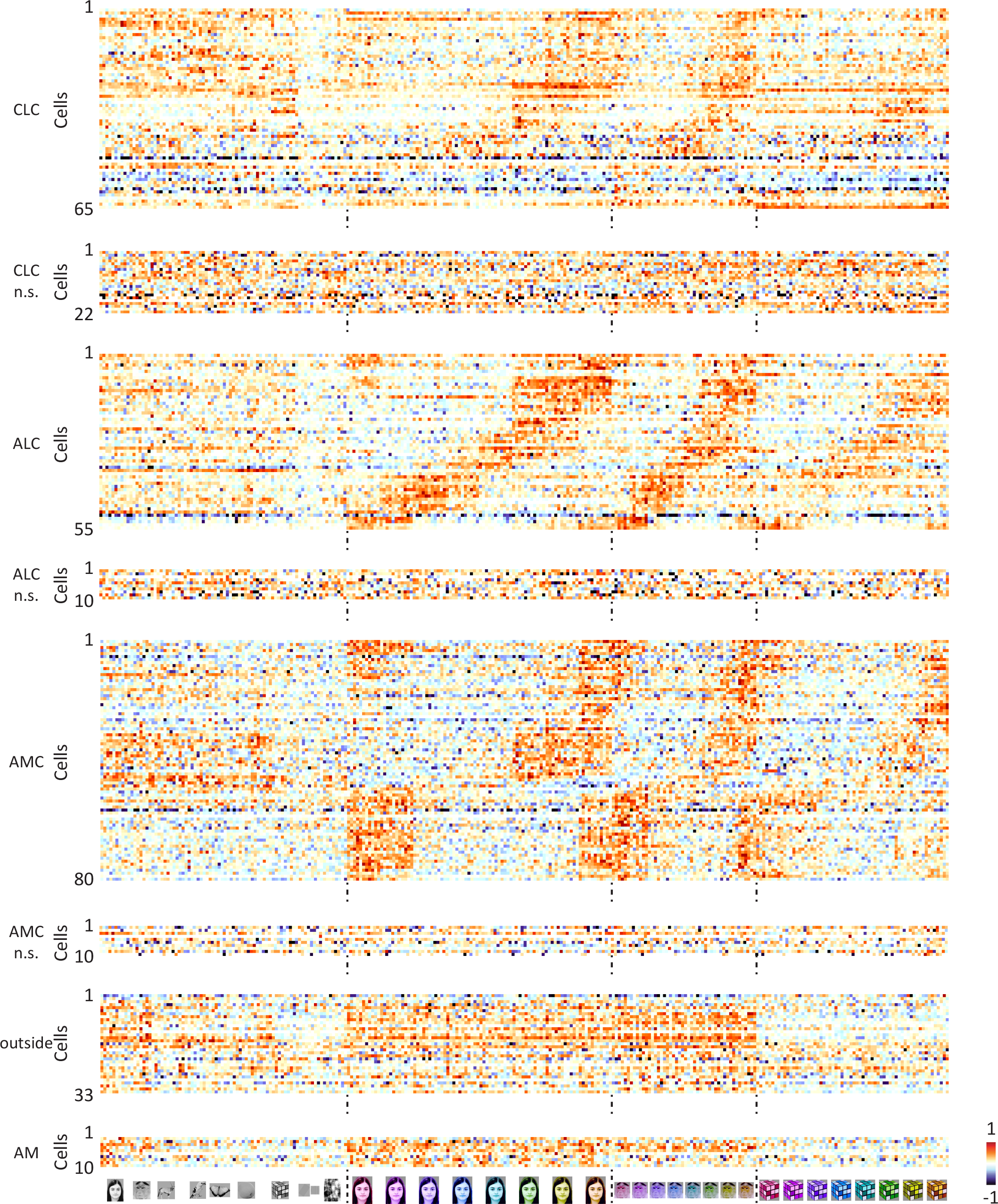
Responses of color patch neurons to the color stimuli. Responses of all neurons in three patches to all grayscale images, colored human faces, monkey faces and magic cubes, sorted according to hue preference of the average response across all 82 stimuli. Color-selective cells and non-selective cells are shown separately. Responses of IT cells outside color patches and cells in face patch AM are also shown. For each cell, baseline was subtracted and the response was normalized.

Responses to the 10 color conditions for all color-selective cells, grouped according to object category, are shown in Fig. 3a. Each row represents one cell; cells in each patch are sorted according to hue preference, computed using responses of each cell to 8 hues averaged across all 82 objects. There is clear consistency of hue tuning across categories, especially for ALC and AMC. Hue consistency was quantified by computing the correlation of hue tuning across categories (Supplementary Fig. 3a; see Supplementary Fig. 3b for correlations between all pairs of categories). We found all neurons demonstrated a positive correlation, with ALC and AMC significantly more consistent than CLC (W(65,55)=3215, p=2*10^−4^ between CLC and ALC; W(65,80)=3473, p=4*10^−7^ between CLC and AMC; W(55,80)=3767, p=0.91 between ALC and AMC, Wilcoxon rank sum test; average correlation value: 0.533 for CLC, 0.686 for ALC, 0.706 for AMC, and 0.119 for outside color patches). Outside the color patches, consistency of hue tuning was much lower (Supplementary Fig. 3a, W(33,200)=803, p=5*10^−17^, Wilcoxon rank sum test).

**Figure 3.**
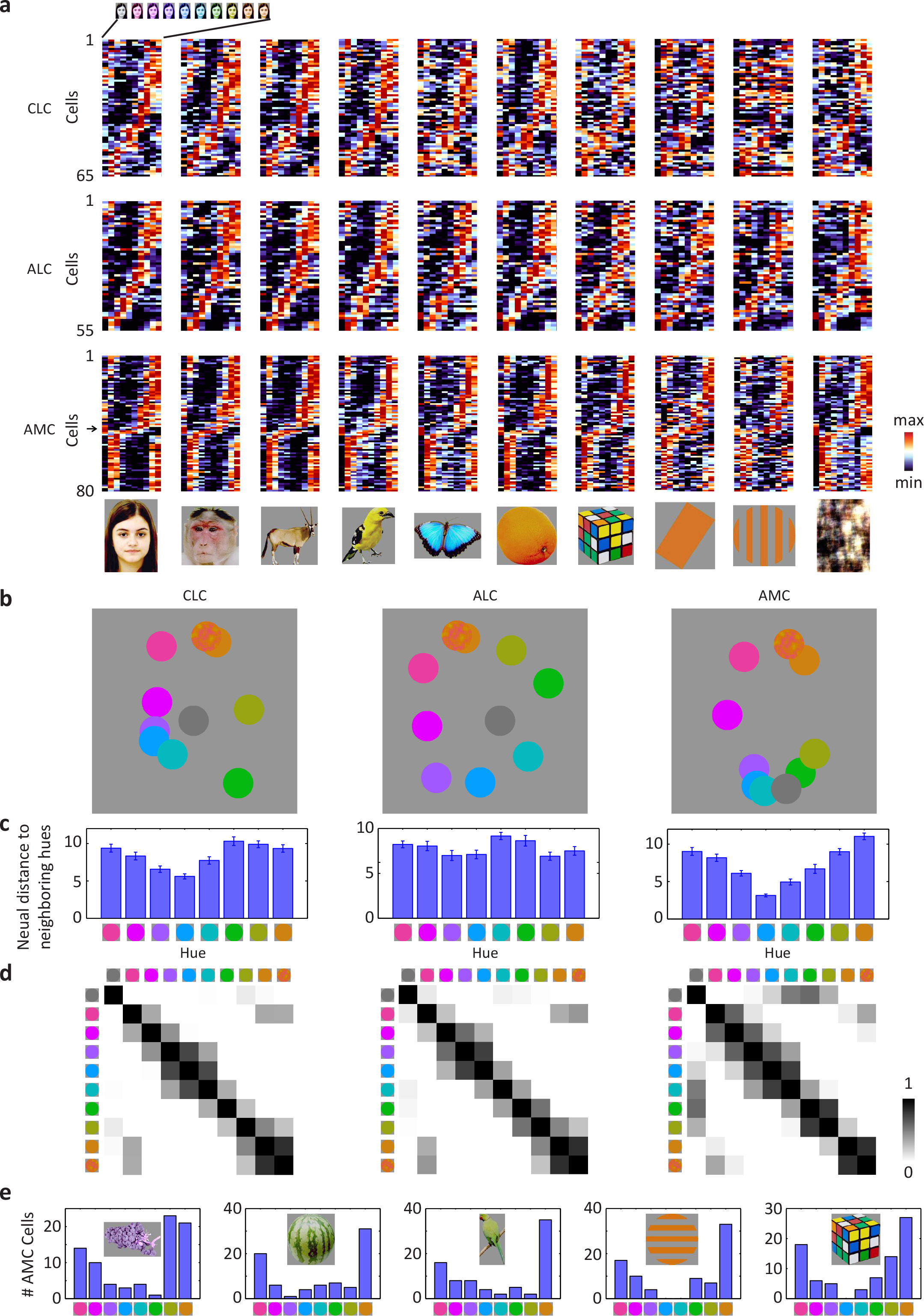
Representation of color by color-selective neurons in color patches. **a,** Responses of all color-selective neurons averaged across stimuli within each category to images of 8 different hues, together with gray (left-most column) and natural color (right most-column), sorted in this same way as Fig. 2. **b,** Neural representation of colors in the activities of color-selective neurons. Shown are two-dimensional plots of the results of multi-dimensional scaling (MDS) analyses conducted for neurons in three color patches. Responses to each color condition were averaged across all mammal images (humans faces, monkey faces, and mammal bodies; these were selected because color tuning was most consistent between these three categories across all three patches, see Supplementary Fig. 3b). Original color is indicated by a disk of mixed color. **c,** Neural distances of each hue to its two neighboring hues, for all three patches, computed using population responses of color-selective cells. Error bars represent s.d. of 20000 iterations of bootstrapping. Inhomogeneity was quantified by computing the ratio between the s.d. of the 8 bars and the mean of the 8 bars: 0.21±0.02 for CLC; 0.12±0.02 for ALC; 0.35±0.02 for AMC (p<0.001 between CLC and AMC; p<0.001 between ALC and AMC; p=0.0103 between CLC and ALC, 20000 iterations of bootstrapping, see Methods). **d,** Population similarity matrices of 10 color conditions in three color patches. A 10*10 matrix of correlation coefficients was computed between responses of all color-selective neurons averaged across objects. **e,** For five different types of objects: grapes, watermelon, birds, gratings and rubik’s cube, the number of AMC cells preferring each of the eight hues was counted. In all five cases, the distribution was significantly different from homogeneity (chi-square test: p<0.001; χ^2^(7)=26.3, 28.6, 31.0, 38.6 and 28.4 respectively).

We observed a difference between hue tuning in CLC/ALC compared to AMC. Most AMC cells preferred red or yellow to other hues, leading to an under-representation of green and blue (Fig. 3a, arrow); in CLC and ALC, different hues were more evenly represented (Fig. 3a; the proportion of cells preferring the 4 hues, from purple to green in CLC, ALC and AMC is 33.9 %, 47.3 % and 8.7 %. Chi-square test, χ^2^(1)=12.61 and p=3*10^−4^ between CLC and AMC; χ^2^(1)=23.11 and p=2*10^−6^ between ALC and AMC; χ^2^(1)=2.22 and p=0.13 between CLC and ALC). Multidimensional scaling (MDS) analyses on population responses revealed an additional difference in hue representation between AMC and the other two patches: In CLC and ALC, the neural representation of all 8 hues is homogeneous, with gray located in the center of eight hues; in AMC, yellow is over-represented, with gray located in the periphery, very close to cyan and green (Fig. 3b). Note that gray is surrounded by the eight hues in the original CIE color space (Fig. 1e), analogous to the population representation in ALC visualized with MDS.

We quantified the transformation in hue representation across CLC, ALC, and AMC by computing the “neural” distance between neighboring hues based on population responses. AMC neurons displayed stronger inhomogeneity in distance between neighbors than the other two patches (Fig. 3c). The difference in color tuning between CLC/ALC and AMC was further confirmed by representation similarity matrices quantifying the correlation between mean population responses to pairs of colors in each of the three patches (Fig. 3d). The correlation between gray and cyan/green is evident in AMC, but almost absent in the other two patches (mean correlation=0.02±0.08 for CLC; −0.02±0.12 for ALC; 0.45±0.09 for AMC; p<0.001 between CLC and AMC; p<0.001 between ALC and AMC; p=0.385 between CLC and ALC, 20000 iterations of bootstrapping). In all three patches, the representation of the original color images was closest to yellow/red (Fig. 3b, d). This could be due to the fact that pixel intensities of the original color images turned out to be tightly distributed around yellow, especially for the faces and bodies (Supplementary Fig. 2b-h). This last fact could also explain why AMC preferred red/yellow hues: it could be biased to represent the color of faces and bodies. However, it’s worth noting that AMC neurons showed preference for red/yellow even for objects that are not naturally red/yellow, e.g. grapes, watermelons, and abstract shapes without any natural color association (Fig. 3e). Thus AMC cells were not simply over-representing correctly-colored objects.

One concern is that the preference for red/yellow observed in AMC could be due to undersampling in our single-unit recordings. We recorded from AMC in ten different penetrations in two monkeys, and results were consistent across both animals. To further address this concern, we performed an fMRI experiment in which we presented red, yellow, blue, and grayscale monkey faces (see below for rationale for showing monkey faces). Contrasting red/yellow versus grayscale monkey faces revealed activation in CLC, ALC, and AMC (Supplementary Fig. 4a). Importantly, activation to red/yellow was significantly stronger than blue in AMC, but not CLC/ALC (Supplementary Fig. 4b). The presence of the bias for red/yellow in the global AMC fMRI signal shows that it is not due to selective sampling.

We also compared sharpness of hue tuning across the three patches, and found evidence for gradual sharpening from CLC to ALC to AMC (Supplementary Fig. 5). Overall, the results so far show that (1) each color patch contains a large population of hue-tuned cells, and (2) a transformation in hue tuning occurs between CLC/ALC, where different hues are uniformly represented, and AMC, where red and yellow are over-represented compared to green/blue.

### Representation of object shape by neurons in color patches

Thus far, we have examined tuning to hue, averaged across object identity. However, since the color patches have a stereotyped location relative to face patches, which represent facial shape, a natural question is: how is object shape represented across color patches? To quantify the representation of object shape, we computed responses to 82 objects averaged across 8 hues. MDS was conducted on population responses of color-selective cells in three patches (Fig. 4a). All three patches displayed clear grouping according to object category. But the amount of information about object shape was very different between the three patches: the accuracy for identifying images, quantified by a nearest neighbor classifier (see Methods), was significantly higher in CLC than ALC across all ten categories (Fig. 4b, p<0.05, 20000 iterations of random sampling with replacement, cell numbers were equalized to 55 for all three patches). Comparing ALC with AMC, identification accuracy was significantly higher in AMC for humans and monkeys (p<0.05) but not other categories (Fig. 4b). Comparing CLC with AMC, accuracy was significantly higher for all categories (p<0.05) except from monkeys (p=0.496). A 2-way ANOVA analysis to test significant interaction between area (2 levels) and category (10 levels) revealed significant interactions for ALC and AMC (F(9)=2.03, p=0.040), CLC and AMC (F(9)=3.36, p=0.001), but not ALC and CLC (F(9)=0.98, p=0.459).

**Figure 4.**
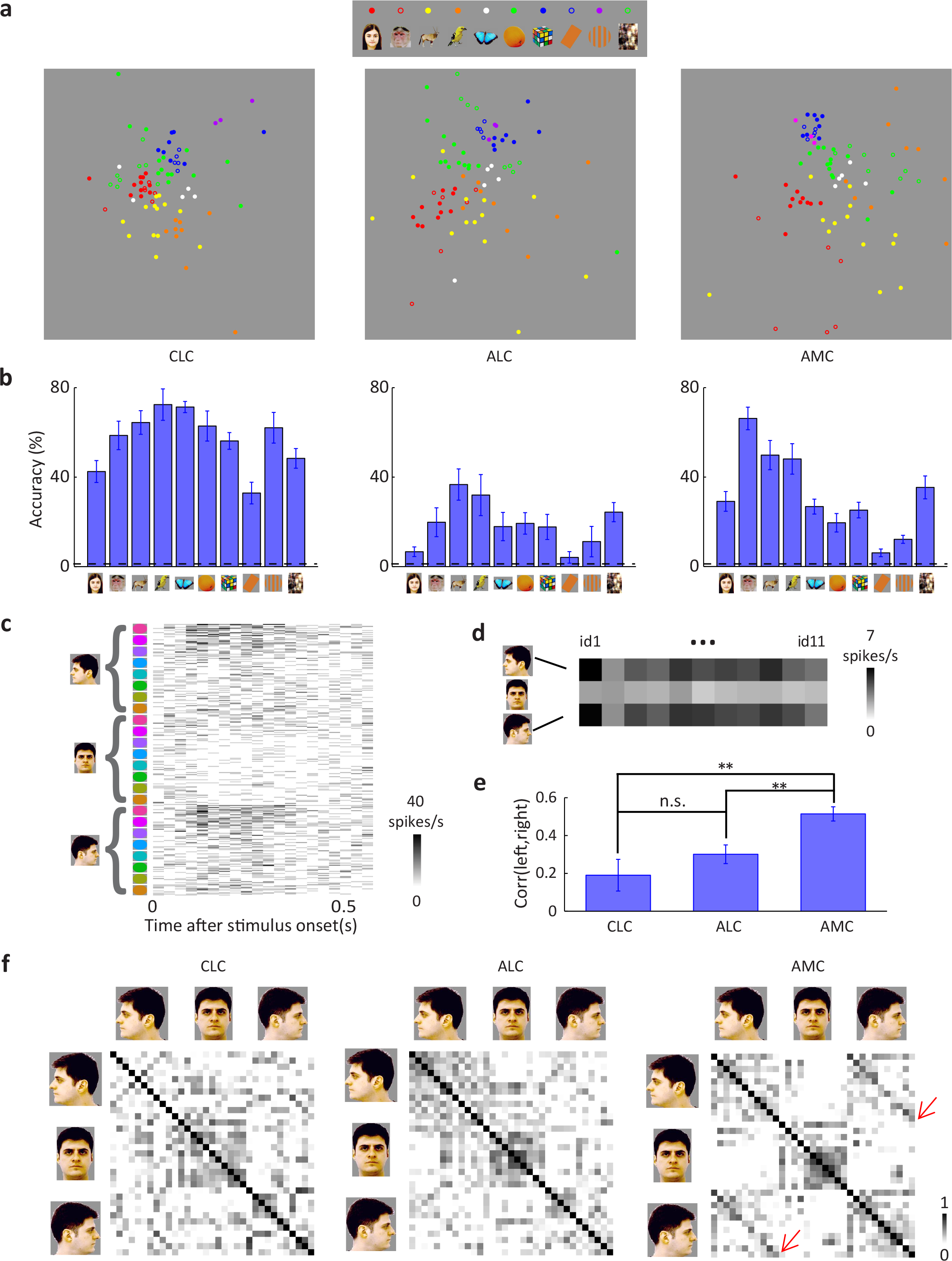
Representation of object shape by color-selective neurons in color patches. **a,** Neural representation of object shapes in the activities of color-selective neurons for three patches, shown as two-dimensional MDS plots. Responses to each object shape were averaged across 8 hues. **b,** Decoding accuracies for identifying one object out of 82 objects based on population responses in three color patches, averaged across object identities within each category (see Methods). Error bars represent s.e. Dashed lines indicate chance level (1/82=1.2%). **c,** Raster plot showing responses of an AMC neuron to colored human faces of 11 identities at three views: frontal, left- and right-profile (Supplementary Fig. 2i). **d,** The same response in (c) averaged across 8 hues, showing strong correlation between left- and right-profiles, but not between frontal and profile views. **e,** Correlation between responses to left and right profile views was computed across identities for each cell. Mean and s.e. of all neurons in three patches are plotted (n=26 CLC cells; n=40 ALC cells; n=44 AMC cells). Student’s t-test was used to determine statistical significance between patches (*=p<0.05, **=p<0.01). **f,** Population similarity matrices of 11 identities*3 views in three color patches. The paradiagonal stripes in AMC indicate high correlation between responses to mirror-symmetric views of the same identity (red arrows).

Could the shape information observed in the color patches be a vestige of low-level shape selectivity present in presumptive inputs to the color patch system, e.g., orientation-tuned cells in area V4? To address this, we compared object shape representations in each color patch with those in two models, AlexNet ^17^ and HMAX ^18^, a model for visual processing in V1-V4. The results show that the shape representations in CLC, ALC and AMC are high-level, consistent with those in other parts of IT cortex (Supplementary Fig. 6a-d). Note that here we only investigated representation of shape independent of hue, thus the similarity between color patches and other regions in IT (Supplementary Fig. 6a) is restricted to shape and does not indicate anything about color representation.

Within the face patch system, the most salient difference between patches is how they represent facial identity across different views, with an increasingly view-invariant representation as one moves anterior ^9^. Does a similar transformation in view-invariant object identity occur in the color patches? We presented facial images of different identities at eight hues and three views: left/ right profiles and frontal (Supplementary Fig. 2i). In AMC, we found cells mirror symmetrically-tuned to views (Fig. 4c, d shows one example cell). The population response showed a correlation between responses to left and right profile views of the same identity (Fig. 4e, f; t(68)=−4.02, p=1*10^−4^ between CLC and AMC; t(82)=−3.48, p=8*10^−4^ between ALC and AMC; t(64)=−1.22, p=0.23 between CLC and ALC, Student’s t-test), similar to anterior face patch AL ^9^. This mirror symmetric view invariance was weaker in CLC and ALC (Fig. 4e, f). This result shows that the representation of facial shape in AMC is not simply inherited from CLC.

### Co-representation of hue and shape by neurons in color patches

The analyses so far have examined color and shape representations in isolation, and revealed that both color and shape information are present in all three color patches. To gain a full picture of information flow across patches, we next examined co-representation of the two information channels across patches.

We first conducted MDS analyses on responses to all human face images in the stimulus set (Fig. 5a). We found that the neural representation of colored faces differed between the three patches: although all three patches showed grouping of images according to hue, this grouping was clearer in ALC and AMC than CLC. Outside the color patches, images were grouped according to identity rather than hue. This difference is illustrated by similarity matrices (Fig. 5b; for full similarity matrices see Supplementary Fig. 7): the 11*11 squares along the diagonal reflecting hue-specific representation are strongest in ALC, followed by AMC, and least clear in CLC; the para-diagonal stripes indicating hue-invariant identity information are only clearly observed in CLC but not the other two patches. Outside the color patches, strong para-diagonal stripes are visible. To quantify relative contributions of hue and identity for each category, we averaged correlation coefficients between population responses to images with the same hue but different identity or same identity with different hue (Fig. 5c). We found comparable amounts of hue and identity information for all 10 categories in CLC (p=0.053, 0.359, 0.203, 0.128, 0.006, 0.025, 0.421, 0.001, 0.287, 0.329, 20000 iterations of bootstrapping), but a clear bias for hue information in ALC (p=0, 0, 0, 0, 0, 0, 0.027, 0, 0, 0.004). AMC was similar to ALC, showing a strong bias for hue, with one prominent exception: for the category of monkey images, hue and identity information were comparable (p=0.34). The presence of hue-invariant identity information about monkeys in AMC is consistent with the superior ability to identify monkeys compared to other objects using AMC population responses (Fig. 5b). The enhanced representation of monkey identities compared to other objects in AMC adds support to our hypothesis that it is biased to represent colored faces and bodies. Outside the color patches, there was significantly more information about identity compared to hue for all the natural image categories (Fig. 5c).

**Figure 5.**
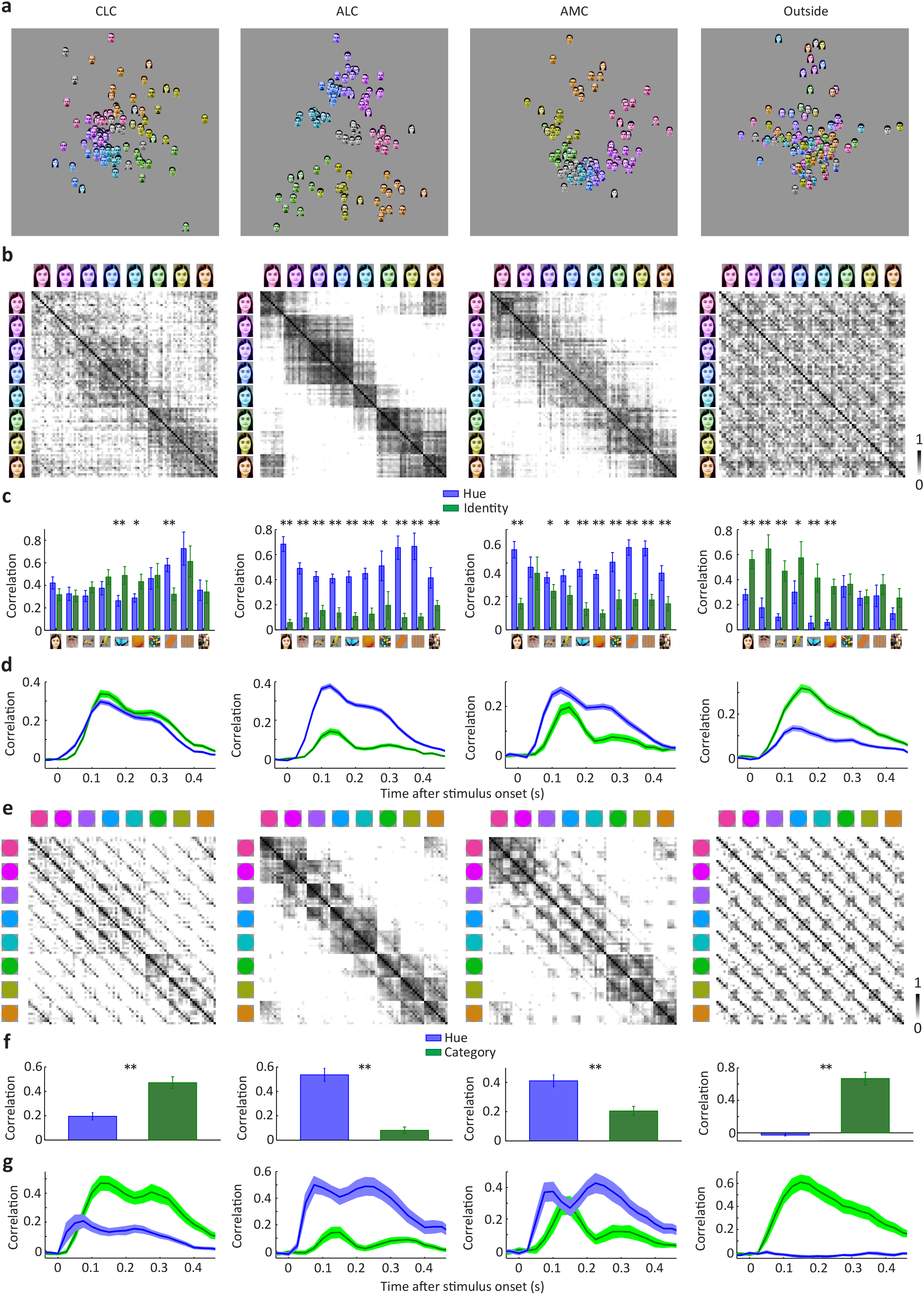
Co-representation of hue and object identity in color patches. **a,** Comparison of MDS plots of responses to all human face images in all three color patches and outside color patches. For clarity, the original natural color images are not shown. In ALC and AMC, images were clearly grouped according to hue, while in CLC this grouping is less clear. Outside the color patches, images were grouped according to identity, but not hue. **b,** Population similarity matrices computed from responses to human face images in three color patches and outside color patches. Correlation coefficients were computed between responses to 11 identities and 8 hues. **c,** Hue information and identity information for images of 10 categories in three color patches and outside color patches. Hue information was quantified as the mean correlation between responses to images with the same hue but different identity within the same category, while identity information was quantified as the mean correlation between responses to images of the same identity with different hues. Note that here we are quantifying shape-invariant hue tuning, which will be affected by both shape tuning and color tuning; in particular, cells with strong shape tuning will show low shape-invariant hue tuning, even if they have perfectly consistent hue tuning across shapes. Error bars represent s.d. of 2000 iterations of bootstrapping. Statistical significance was determined between hue and identity information for each category in three color patches and outside color patches (*=p<0.05, **=p<0.01). **d,** Amplitude of hue and identity information for three patches and outside color patches, computed over a 50 ms sliding time window, were averaged across all 10 categories. Shaded regions indicate s.d. estimated by 2000 iterations of bootstrapping. **e,** Co-representation of hue and category in all three patches and outside color patches. Responses of each cell were averaged across different identities within a category. A matrix of correlation coefficients was computed between responses to 10 categories* 8 hues. **f and g,** Same as (c) and (d), but for hue and category information quantified by matrices in (e), **=p<0.01.

We further analyzed the co-representation of hue information and category information using responses averaged across all identities within one category. This revealed a bias for category information in CLC and IT regions outside the color patches, and hue information in ALC and AMC (Fig. 5e, f). Examination of the time course of color information and identity information in all three patches revealed that color was always faster than identity, even in CLC where identity information was slightly stronger at the peak (Fig. 5d, g). To quantify the difference in temporal dynamics, we defined latency as the first time point hue/shape information differed significantly from baseline (p<0.01, 20000 iterations of bootstrapping, note that the each time point t indicates a time window [t-25 ms, t+25 ms]). In all cases, color was faster than shape (shape identity vs. color: i.d.=50 ms and color=25 ms for CLC, i.d.=75 ms and color=50 ms for ALC, i.d.=75 ms and color=50 ms for AMC; shape category vs. color: category=50 ms and color=25 ms for CLC, category=75 ms and color=25 ms for ALC, category=75 ms and color=50 ms for AMC).

Previous studies on object representation in IT from other labs employed linear classifiers to quantify the amount of “linearly” decodable information from population response of neurons ^14 19^. This provides a useful way to measure whether a particular dimension of information coded by a certain brain area is “untangled” from other dimensions. We applied the same method to our data to quantify the amount of shape-invariant hue information and hue-invariant shape information. Consistent with our analyses with similarity matrices, we found that shape-invariant hue information was higher in anterior color patches than in CLC, while hue-invariant shape information showed the opposite trend (Fig. 6). Thus suggests that hue information does indeed become untangled from shape information along the color patch pathway.

**Figure 6.**
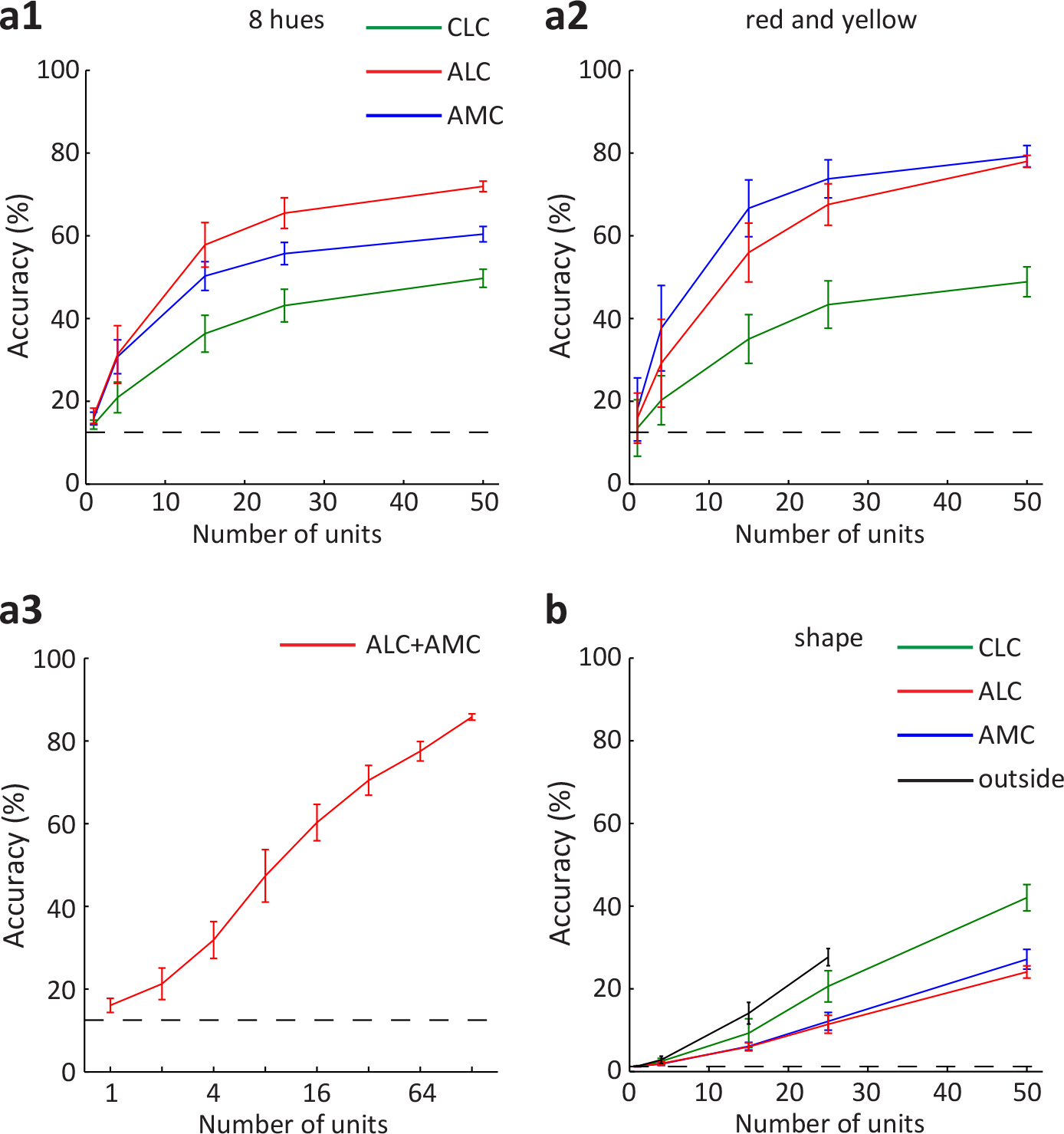
Decoding shape-invariant color and color-invariant shape from color patches. **a1,** Svm models were trained to classify hues independent of shape. Population response of different number of randomly selected units was used as the input to the model. Half of the trials were used for training and the rest half for cross-validation. Shape-invariant hue could be significantly better decoded by AMC and ALC population than than CLC (for 50 units, p<0.01 for both comparisons, 2000 iterations of bootstrapping). Furthermore, ALC shows better overall decoding than AMC (p=0.013). Dashed line indicates chance level (1/8=12.5%). Results are averages across 2000 iterations of random sampling. Errorbars represent s.d. **a2,** similar to (a1), but only quantifies decoding accuracy for two hue categories: red and yellow. Decoding based on AMC is better than ALC, but not significant (p=0.185). **a3,** similar to (a1), but for a combined population of anterior color patch neurons. **b,** similar to (a1), but for hue-invariant shape decoding. CLC is significantly better than ALC and AMC (for 50 units, p<0.01 between CLC and ALC, p=0.028 between CLC and AMC). Furthermore, neurons outside color patches shows better performance than color patch neurons, but only significantly better than ALC and AMC (For 25 units, p<0.01 between outside and ALC, p=0.019 between outside and AMC and p=0.186 between outside and CLC). Dashed line indicates chance level (1/82=1.2%).

### Co-representation of hue and object shape at the single-cell level

So far, we have shown that all three patches contain information about both object shape and hue. What is the relative contribution of these two variables within single cells? Furthermore, what integration rule is used by single cells in color patches to combine hue and shape information? To answer these two questions, we performed a 2-way ANOVA analysis on single cell responses (Fig. 7a), with 82 levels of shape and 8 levels of hue. A scatter plot of explained variance due to hue versus that due to shape revealed an inverse relationship between the two, as expected (Fig. 7b1). Consistent with previous population analyses, AMC and ALC neurons are more biased to hue than CLC neurons (Fig. 7b2): ALC and AMC are significantly more hue-biased than CLC (t(118)=−7.61, p=7*10^−12^ and t(143)=−9.0, p=2*10^−15^ respectively, Student’s t-test); CLC is significantly more hue-biased than outside (t(96)=−6.1, p=3*10^−8^); ALC and AMC are not significantly different (t(133)=0.2, p=0.85). However, it is worth noting that all three color patches did carry a significant amount of shape information. In CLC, the mean amount of variance accounted for by shape was 37.4%. In ALC, the mean amount of variance accounted for by shape was 18.8%, while in AMC, it was 20.2% (Fig. 7b2; note that given noise in the data, the relative contribution from shape is over-estimated, since there are more parameters for shape (82) than for hue (8)). We performed similar analysis, but using 10 coarse shape categories or shape-within-categories (on average 8.2 shapes in each category) to define the shape variables. We found that in these two cases, AMC and ALC neurons, but not CLC neurons, were clearly hue biased (Fig. 7b3-b4). Comparing four regions for coarse shape categories: ALC and AMC are significantly more hue-biased than CLC (t(118)=−4.5, p=1*10^−5^ and t(143)=−4.6, p=8*10^−6^ respectively, Student’s t-test); CLC is significantly more hue-biased than outside (t(96)=−7.9, p=4*10^−12^); ALC and AMC are not significantly different (t(133)=0.6, p=0.5). Comparing four regions for shape-within-categories: ALC and AMC are significantly more hue-biased than CLC (t(118)=9.0, p=3*10^−15^ and t(143)=9.5, p=3*10^−17^ respectively, Student’s t-test); CLC is significantly more hue-biased than outside (t(96)=−6.5, p=3*10^−9^); ALC and AMC are not significantly different (t(133)=−0.8, p=0.4). As expected, all color patch neurons show a significant main effect for hue (ANOVA, p<0.001), while most color patch neurons (with the exception of 2 ALC neurons and 1 AMC neuron) showed a significant main effect for shape (Fig. 7b5). Finally, we found a large portion of color patch neurons showed a significant interaction between hue and shape (41/65 (=63%) CLC cells, 40/55 (=73%) ALC cells, 47/80 (=59%) AMC cells, and 0/33 cells outside color patches, Fig. 7b6).

**Figure 7.**
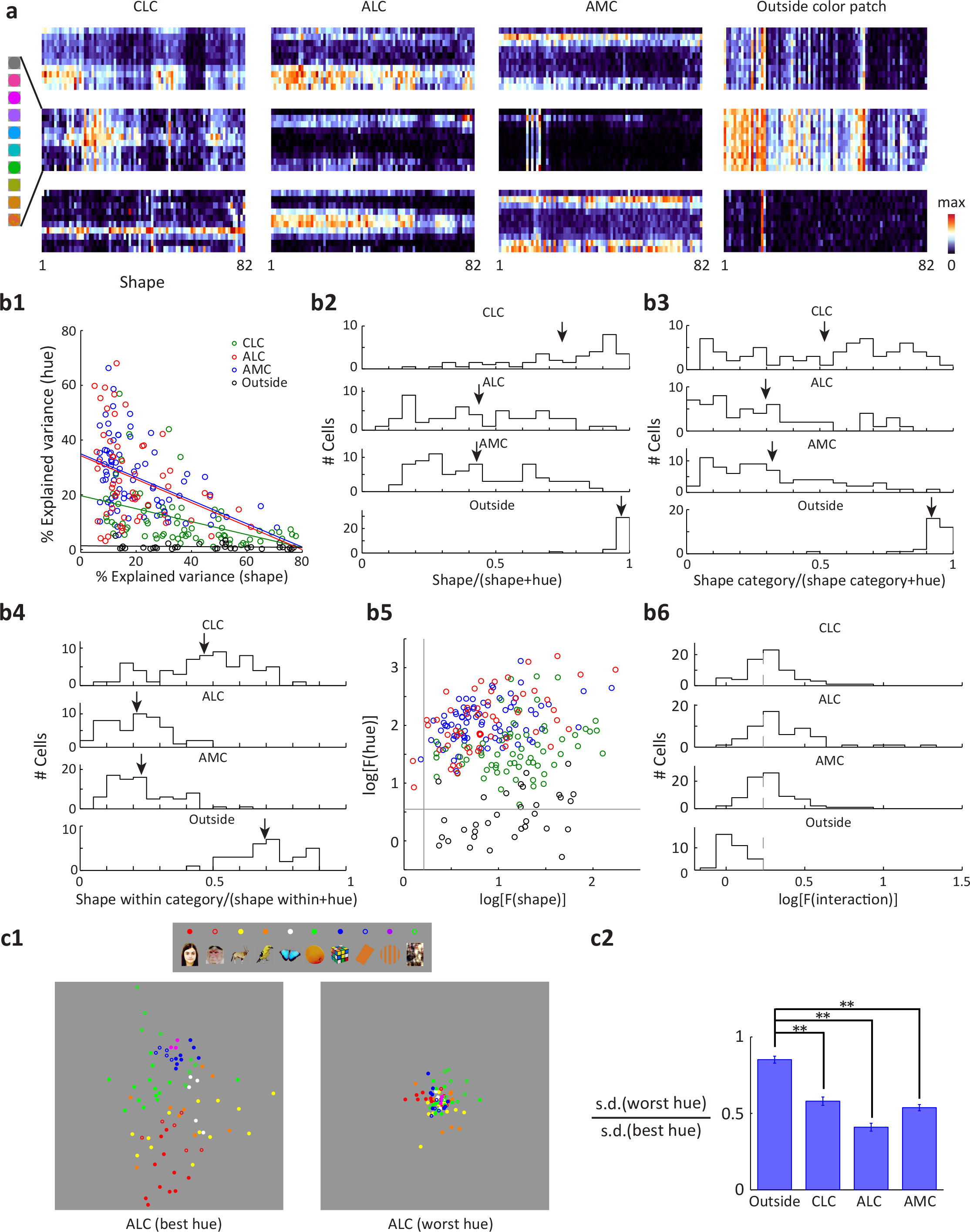
Analysis of single-cell responses. **a,** Responses of 12 example neurons to the full stimulus set. Each row represents one color condition, and each column represents one object shape. **b,** 2-way ANOVA analysis examining main effects of shape and hue, as well as interactions. 2-way ANOVA analysis with 8 levels of hue and 82 levels of shape was performed on responses of each individual neuron. **b1,** Relationship between explained variances by two main effects for all neurons. Lines represent linear fits to cells in each patch. **b2,** Distribution of shape preference in all three patches and outside the color patches, defined by the explained variance by shape divided by the sum of both main effects. Arrows indicate population averages. **b3,** similar to (b2), but using only coarse shape categories as shape variables. **b4,** similar to (b2), but using only fine shapes within each shape category as shape variables. ANOVA analysis was carried out for each shape category independently, with 8 levels of hue and n levels of shape (n=number of shapes within this shape category). For each neuron, shape preference was computed and averaged across categories. **b5,** For 2-way ANOVA with 8 levels of hue and 82 levels of shape, F-values for both main effects are plotted against each other in log-scale. Gray lines indicate significance level (p=0.001). **b6**, Distribution of F-values for the interaction between hue and shape. Gray dashed lines indicate significance level (p=0.001). **c1,** For each cell in ALC, we determined the best hue and the worst hue based on responses to 8 hues averaged across 82 object shapes. MDS analyses were conducted on shape responses for the best hue or the worst hue of each cell. Two MDS plots are shown at the same scale. **c2,** For each cell, the ratio between standard deviations of shape responses at the worst hue and the best hue was computed. If the cells were linearly adding hue and shape, the two standard deviations should be identical. Therefore the ratio between these two reflects the extent of nonlinearity in the interaction of hue and shape. **=p<0.01, Student’s t-test.

The nonlinear interaction between shape and hue is further supported by comparison of MDS analysis of shape responses at the best and worst hues. If cells were linearly adding the two variables, then the MDS plots at the two hues should be identical; however, we found that shape representation at the worst hue was compressed compared to that at the best hue (Fig. 7c1, c2; t(96)=6.4 and p=7*10^−9^ between outside vs. CLC; t(86)=11.8 and p=1*10^−15^ between outside and ALC; t(111)=9.0 and p=6*10^−15^ between outside and AMC, Student’s t-test). Overall, these results suggest that cells in color patches are not simply summing shape and color inputs, but are nonlinearly combining the two.

The presence of nonlinear interaction between shape and hue raises the question whether the main effects for hue and shape are real, i.e. the main effects may appear only in some conditions but not at all in others. To address this, for each cell in each patch, we quantified hue tuning for each shape by computing the variance of responses across hue to the shape, and selected 27 shapes with the best and worst hue tuning. Similarly, we selected 3 hues with the best and worst shape tuning for each cell. We computed the correlation between hue tuning for the “best” and the “worst” shapes, and did the same for shape tuning. We found all cells except 1 ALC cell showed positive correlation between hue tuning for the “best” and the “worst” shapes, and most cells (65/65 CLC cells, 51/55 ALC cells, 71/80 AMC cells) showed positive correlation between shape tuning for the “best” and the “worst” hues. Furthermore, we performed ANOVA on the “worst” shapes/hues, and a high percentage of cells showed significant (p<0.001) main effects for hue (34/65 CLC cells, 42/55 ALC cells, 69/80 AMC cells)/shape (64/65 CLC cells, 47/55 ALC cells, 69/80 AMC cells).We note that different selection criterion was applied to color patches and neurons outside the color patches in the above two analyses (Figs. 5 and 7): while only color selective cells were used for color patches, all cells were pooled for “outside”. However, all the results comparing color patches to outside the color patches remained consistent when we pooled all cells in color patches.

## Discussion

In this study, we found that three macaque color patches, CLC, ALC, and AMC, all encode significant information about both hue and object identity. Two clear transformations occur across the three patches. The first transformation, from CLC to ALC, reduces information about object identity. The second transformation, from ALC to AMC, mainly affects representation of hue: color space is represented in a dramatically distorted way in AMC, with over-representation of yellow and red, the natural colors of mammal faces and bodies. Furthermore, AMC develops an expanded representation of primate faces compared to other categories, displaying hue-invariant representation of monkey identity.

Our study broadens our conception of the function of IT cortex. A generally accepted notion is that the purpose of IT is to represent object identity invariant to accidental changes, and this is achieved through a hierarchy culminating in cells in anterior IT tuned to object identity invariant to accidental changes ^9,18,20^. For example, in the face patches, tuning to facial identity becomes more invariant to view going from ML/MF to AL to AM ^9^. Applying this principle to colored objects, one might expect to find a sequence of areas tuned to object hue and shape combinations that show increasing invariance to accidental transformations in view, lighting, etc. (Fig. 8b). Instead, we found the existence of a specialized network in which shape information *decreases* along the IT hierarchy, while hue information *increases* (Fig. 8a). Since shape is one of the strongest cues to object identity, this calls into question the current picture of IT as a monolithic hierarchical feedforward network for computing object identity ^20^. IT appears to generate an array of high-level representations of objects that can facilitate different object-related tasks, including the fundamental task of identifying the color of an object.

**Figure 8.**
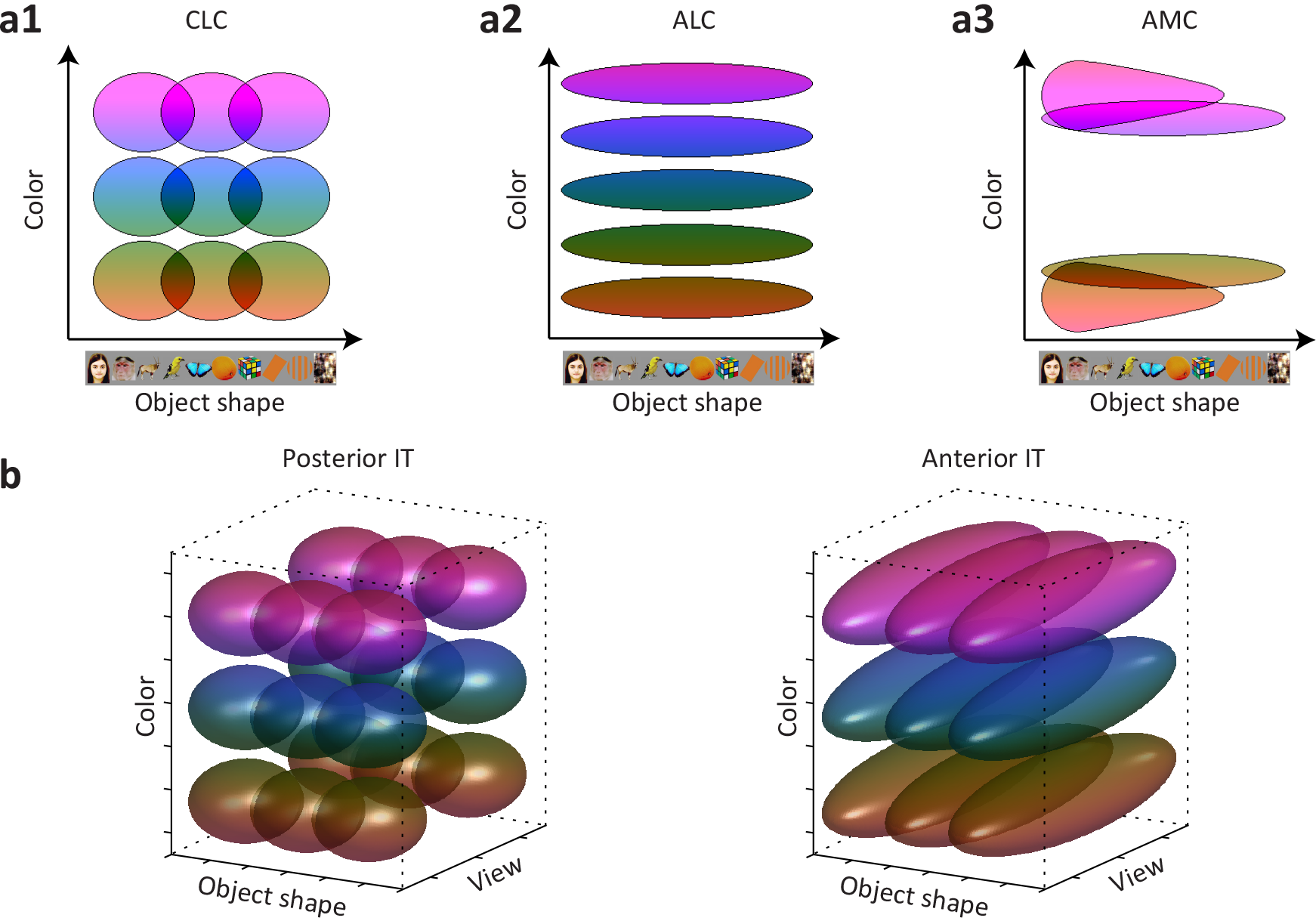
Extending the conventional view of IT: a theory of color processing in IT. **a,** Schematic summary of the co-representation of hue and object shape in three color patches. Here, each oval represents the receptive field of one “idealized” color neuron in the 2-d object space spanned by hue and object shape. **b,** Conventional view of IT predicts that the major transformation of colored-object representation from posterior to anterior IT is the generation of invariance to accidental changes (e.g., view). Here each ellipsoid represents the receptive field of one “idealized” neuron in the 3-d object space spanned by hue, shape, and view. For both posterior and anterior IT, two dimensional slices at a fixed “view” should look the same as the schematic for CLC (a1).

What is the role of AMC? Why should there exist an area with neurons that represent hue irrespective of shape, but mainly for red and yellowish things, but then also the shape of faces irrespective of hue? At first glance, this seems unparsimonious. One possible explanation is that AMC provides an important intermediate link between a multi-purpose shape-invariant hue representation and a representation specialized for the color of animal faces/bodies. The diversity in responses to different faces in AMC could compensate for the relative homogeneity in their responses to hues, exploiting the coding space previously occupied (in ALC) by bluish hues. This would facilitate the wiring of classifiers trained to identify faces using both shape and color cues. Future experiments exploring responses in color patches during performance of active tasks including face and color categorization may shed further light on the functional role of each patch.

Few previous studies have examined color tuning in IT cortex. Most have reported aggregate statistics based on random sampling of IT cortex ^21–25^, and have come to conflicting conclusions regarding the prevalence of hue-tuned cells, from 15% ^21^ to 69% ^22^, as well as the extent to which shape and hue interact in single cells; Komatsu and Ideura ^22^ reported no interaction, while Edwards et al. ^23^ reported strong nonlinear interaction. One pioneering study of color transformations in IT reported two clusters of color-selective cells in IT cortex, one in posterior and one in anterior IT, and showed a difference between these two regions in their color tuning as a function of luminance ^12^. Our study shows that there are at least three clusters of color-selective cells in IT that are strongly anatomically connected. Most importantly, our study demonstrates the importance of (1) studying the *co-representation* of color and object shape within each color patch, and (2) studying multiple patches within the IT color network using the same stimuli. Only by taking both of these steps, could we reveal the transformations in the brain’s representation of colored objects for the first time.

## Methods

### Color Patch Localization

All procedures conformed to local and US National Institutes of Health guidelines, including the US National Institutes of Health Guide for Care and Use of Laboratory Animals. All experiments were performed with the approval of the Caltech Institutional Animal Care and Use Committee (IACUC).

Four male rhesus macaques were trained to maintain fixation on a small spot for juice reward (three of the animals were used for color patch recordings, while the fourth animal was used solely for control recordings outside color patches). Monkeys were scanned in a 3T TIM (Siemens, Munich, Germany) magnet equipped with AC88 gradient insert while passively viewing images on a screen. Feraheme contrast agent was injected to improve signal/noise ratio. Color patches were determined by identifying regions responding significantly more to moving equiluminant red/green color gratings (2.9 cycles per degree, drifting 0.75 cycles per s) than moving black-white gratings, the same stimuli as a previous study ^5^, and were confirmed across multiple independent scan sessions. In monkey M1, color patches CLC, ALC and AMC were found bilaterally. In monkey M2, color patches CLC and ALC were found bilaterally; color patches AMC and AFC were found unilaterally in the left hemisphere (AMC was found by microstimulating ALC in the same hemisphere, see below). In monkey M3, color patches CLC and ALC were found bilaterally; color patch AFC was found unilaterally in the left hemisphere.

### Microstimulation

To reveal the anatomical connectivity of color patches and to localize the most anterior color patch AMC in monkey M2, we stimulated ALC ^15^. The stimulation protocol followed a block design. We normally interleaved 9 blocks of fixation-only with 8 blocks of fixation plus electrical microstimulation; we always started and ended with a fixation-only block. During microstimulation blocks we applied one pulse train per second, lasting 200 ms with a pulse frequency of 300 Hz. Bipolar current pulses were charge balanced, with a phase duration of 300 μs and a distance between the two phases of 150 μs. We used a current amplitude of 300 μA. Stimulation pulses were delivered with a computer-triggered pulse generator (S88X; Grass Technologies) connected to a stimulus isolator (A365; World Precision Instruments),which interfaced with different and indifferent electrodes through a coaxial cable. All stimulus generation equipment was stored in the scanner control room; the coaxial cable was passed through a wave guide into the scanner room. We performed electrophysiological recording at the site of stimulation immediately prior to stimulation, to confirm correct electrode placement, as revealed by a high number of hue-selective units.

### Single-unit recording

Tungsten electrodes (18–20 Mohm at 1 kHz, FHC) were back loaded into plastic guide tubes. Guide tubes length was set to reach approximately 3–5 mm below the dura surface. The electrode was advanced slowly with a manual advancer (Narishige Scientific Instrument, Tokyo, Japan). Neural signals were amplified and extracellular action potentials were isolated using the box method in an on-line spike sorting system (Plexon, Dallas, TX, USA). Spikes were sampled at 40 kHz. All spike data was resorted with off-line spike sorting clustering algorithms (Plexon). Only well-isolated units were considered for further analysis. We targeted patches CLC (n=35; right hemisphere) and AMC (n=24; right hemisphere) in monkey M1, patches CLC (n=52; right hemisphere), ALC (n=43; left hemisphere) and AMC (n=66; left hemisphere) in monkey M2, and patch ALC (n=22; right hemisphere) in monkey M3 for single-unit recordings. In addition, we targeted face patch AM (n = 10; right hemisphere) and a region of anterior IT on the ventral bank of the inferotemporal gyrus outside the color patches in M1 (n=10; right hemisphere), and a region of middle IT on the ventral bank of superior temporal sulcus outside the color patches in M4 (n = 13; left hemisphere). Electrodes were lowered through custom angled grids that allowed us to reach the desired targets; custom software was used to design the grids and plan the electrode trajectories ^26^. For each patch, results were qualitatively the same across different monkeys and therefore were pooled together for population analyses. Multiple different tracks were used to target each patch; in particular, for AMC, we designed ten distinct approach angles using different grids to ensure even sampling.

### Behavioral Task and Visual Stimuli

Monkeys were head fixed and passively viewed the screen in a dark room. Stimuli were presented on a CRT monitor (DELL P1130). The intensity of the screen was measured using a colorimeter (PR650, Photo Research) and linearized for visual stimulation. Screen size covered 27.7*36.9 visual degrees and stimulus size spanned 5.7 degrees. The fixation spot size was 0.2 degrees in diameter. Images were presented in random order using custom software. Eye position was monitored using an infrared eye tracking system (ISCAN). Juice reward was delivered every 2–4 s if fixation was properly maintained.

For visual stimulation, all images were presented for 200 ms interleaved by 200 ms of a gray screen. Each image was presented 7–10 times to obtain reliable firing rate statistics. In this study, 2 different stimulus sets were used:

1. A set of 82 images of 10 different categories, varied in 8 different hues. Original images and grayscale images with the same luminance profile were also presented (Supplementary Fig. 2a, for details see below).
2. A set of 33 human face images (Supplementary Fig. 2i), 3 different views of 11 identities, varied in 8 different hues.

### Color stimulus generation

For our color stimuli, we started with a set of 55 object images collected from the internet, and 11 frontal human faces from an on-line database (FEI face database: http://fei.edu.br/~cet/facedatabase.html). We transformed the color of each image in the following way: For a given pixel with RGB value (r,g,b), its chromaticity coordinates and luminance (u,v,L, CIE 1960) were estimated by first computing the frequency spectrum of the pixel by summing the frequency spectra for r, g, and b, each measured separately using a spectrophotometer (PR650, Photo Research), and then converted into chromaticity coordinates (http://www.cvrl.org). The mean luminance of each image was equalized to the background (2.9 cd/m^2^). We then computed the distance of chromatic coordinates of each pixel to “white” (u=0.2105, v=0.3158, filled circle in Fig. 1e). Eight different colors with the same distance but varying angles (open circles in Fig. 1e, starting from 0°, going clockwise at 45° step) were then computed and converted back into an RGB value keeping the luminance (L) unchanged. Repeating this for every pixel resulted in 8 images of pure hues (Fig. 1e right). A grayscale image with same luminance, but “white” chromatic coordinate, was also generated. The stimulus set included simple geometric patterns (8^th^ and 9^th^ row in Supplementary Fig. 2a). For these images, we set the hue of the original image to orange, with mean luminance equal to background. We also included a category of phase scrambled images (last row in Supplementary Fig. 2a). Eight images from the first nine categories were randomly selected and phase-scrambled, keeping the relative phase between different cone components constant.

The other stimulus set was generated in the same way, but using only face images with different head orientations (left profile, frontal and right profile, Supplementary Fig. 2i).

### Color selectivity

For each repetition of the stimulus set, responses to each of the eight hues were averaged across 82 objects. Classical analysis of variance (ANOVA) was performed to test the statistical significance of the differences among eight hue groups, each group containing multiple repetitions of the stimulus set. Only significantly tuned cells (p<0.001) were used for further analysis.

### Multi-dimensional scaling

The number of spikes in a time window of 50-350 ms after stimulus onset was counted for each stimulus. The responses of each cell to all stimulus conditions were normalized to 0 mean and unit variance. To study the neural representation of a single feature, such as hue (Fig. 3b) or object shape (Fig. 4a), responses were averaged across the irrelevant feature (shape in Fig. 3b and hue in Fig. 4a). Classical multi-dimensional scaling was performed on the population responses in each patch, using a Euclidean distance metric and the MATLAB command cmdscale.

### Neural distance

The responses of each cell to all stimulus conditions were normalized to 0 mean and unit variance. Euclidean distance between the normalized population responses to two stimulus conditions was used to quantify the “neural” distance between these two conditions (Fig. 3c).

### Similarity matrix

Based on the same normalized population response, an n × n similarity matrix of correlation coefficients was computed between the population response vectors (across all color-selective cells, averaged over stimulus repeats) to each of the n interested conditions.

### Decoding analysis

To quantify the amount of information about object shape in all three patches, we trained a nearest neighbor classifier: the population response for one particular object averaged across hues in two thirds of the trials was used to define a “template” response for that object. For testing, the population response to one image averaged across the remaining one third of the trials (but not across hues) was compared to each of the 82 “templates”, and the object “template” with minimal distance to actual response was defined as the output of the classifier.

We also employed SVM decoding models as in previous papers ^14,19^. In brief, we randomly selected a number of units from each area, and trained an SVM model for each selection to decode hue information independent of shape or shape information independent of hue using “one vs. rest” approach. We used half the trials to train the SVM model and the remaining half to validate the model. The results shown are validated accuracies.

### Convolutional neural network modeling

To investigate shape representation in three color patches, we loaded 82 objects with 8 different hues into two pre-trained neural networks: 1) a matlab implementation of Alexnet ^17^: http://www.vlfeat.org/matconvnet/pretrained/. This network contains 21 layers: 1^st^, 5^th^, 9^th^, 11^th^ and 13^th^ layers are the outputs of convolution units; 2^nd^, 6^th^, 10^th^, 12^th^ and 14^th^, 17^th^ and 19^th^ layers are the results of rectification; 3^rd^ and 7^th^ layers are the results of normalization; 4^th^, 8^th^, 15^th^ layers are the results of max pooling; 16^th^, 18^th^ and 20^th^ layers are fully connected layers; 21^st^ layer is the output layer (softmax). This network has been pre-trained to identify a thousand objects. 2) a matlab implementation of HMAX model ^18^: http://maxlab.neuro.georgetown.edu/hmax.html#code. This network implements the basic architecture of the HMAX model (S1, C1, S2, C2) and has been pretrained with a set of random natural images. Activations of each unit to each object were averaged across hues to analyze the representation for shape alone. For the case of the HMAX model, since the network only allows grayscale images as input, we presented the grayscale version of the 82 objects to the network to analyze shape representation.

### Population statistics

To determine statistical significance for parameters estimated using population responses, such as correlation, a bootstrap method was employed: neurons were randomly sampled from the population with replacement; 20000 bootstrap samples with equal numbers of neurons were created. A population statistic was computed for each bootstrap sample. The P-value of the null hypothesis was determined by comparing population statistics from 20000 iterations of bootstrapping.

## Acknowledgments

This work was supported by the Howard Hughes Medical Institute, the Tianqiao and Chrissy Chen Institute for Neuroscience at Caltech, and the Swartz Foundation (fellowship to LC). We thank Nicole Schweers for technical support, members of the Tsao lab, Margaret Livingstone, and Simon Kornblith for critical comments.

## Author contributions

L.C. and D.Y.T. designed the experiments, interpreted the data, and wrote the paper. L.C. and P.B. conducted the experiments and analyzed the data.

## Additional information

**Competing interests:** The authors declare no competing financial interests.

